# An Interplay of Phytoplankton Donor Species and Transformation of Released Compounds over Time Defines Bacterial Communities Following Phytoplankton DOMp Pulses

**DOI:** 10.1101/2022.12.23.521850

**Authors:** Falk Eigemann, Eyal Rahav, Hans-Peter Grossart, Dikla Aharonovich, Maren Voss, Daniel Sher

**Author notes:** Corresponding author: Falk Eigemann, Straße des 17 Juni 135, KF 305, 10623 Berlin, Germany.

## Abstract

Phytoplankton-bacteria interactions are stimulated by phytoplankton-released dissolved organic matter (DOMp). Two factors that shape the accompanying bacterial community are i) the “donor” phytoplankton species, defining the initial composition of released DOMp, and ii) the DOMp transformation over time. We added phytoplankton DOM from two globally abundant species - the diatom *Skeletonema marinoi* and the cyanobacterium *Prochlorococcus* MIT9312 - to natural bacterial communities in the Eastern Mediterranean and determined the bacterial responses over a time-course of 72 h in terms of cell numbers, bacterial production (BP), alkaline phosphatase activity (APA), and changes in active bacterial community compositions based on rRNA amplicon sequencing. Both DOMp types were demonstrated to serve the bacterial community as carbon and potentially phosphorus source. Diatom-derived DOM induced higher BP and lower APA compared to cyanobacterium DOM after 24 h, but not after 48 and 72 h of incubation, and also maintained higher Shannon diversities over the course of the experiment, indicating a better bacterial accessibility and broader disposability of diatom derived DOM. Bacterial communities significantly differed between DOMp types as well as different incubation times, pointing to a certain bacterial specificity for the DOMp donor as well as a successive utilization of phytoplankton DOM by different bacterial taxa. The highest differences in bacterial community composition with DOMp types occurred shortly after additions, suggesting a high specificity towards highly bioavailable DOMp compounds. We conclude that phytoplankton associated bacterial communities are strongly shaped by an interplay between phytoplankton donor and the transformation of its released DOMp over time.

**IMPORTANCE:** Phytoplankton-bacteria interactions maintain biogeochemical cycles of global importance. Phytoplankton photosynthetically fix carbon dioxide and subsequently release the synthesized compounds as dissolved organic matter (DOMp), which becomes processed and recycled by heterotrophic bacteria. Yet, the combined effect of the phytoplankton donor species and time-dependent transformation of DOMp compounds on the accessibility to the bacterial community has not been explored until now. The diatom *Skeletonema marinoi* and the cyanobacterium *Prochlorococcus* MIT9312 are globally important phytoplankton species, and our study revealed that DOMp of both species was selectively incorporated by the bacterial community. The donor species had the highest impact shortly after DOMp appropriation, and its effect diminished over time. Our results improve the understanding of biogeochemical cycles between phytoplankton and bacteria, and solve yet unresolved questions of phytoplankton-bacteria interactions.

## INTRODUCTION

Phytoplankton is responsible for ∼50% of the photosynthesis on Earth (1), but a high fraction of the afore photosynthetically fixed carbon is liberated shortly after the buildup (2) in the form of dissolved organic matter (DOMp). The released DOMp serves marine heterotrophic bacteria as energy and carbon (C) source (3), but also contains nutrients such as nitrogen (N) and phosphorus (P), which are likewise consumed and recycled by bacteria (4-7). Hence, interactions between phytoplankton and bacteria are crucial for the functioning of aquatic systems (8-13) and shape biogeochemical cycles at a global scale (2, 11, 14). Different phytoplankton groups and species have different biochemical compositions (15-17), and release different types of DOMp (8, 18-21). Diatoms, for instance, produce especially polysaccharide-rich DOM (22), and diatoms and cyanobacteria own different ratios of specific amino acids in their released DOMp (23, 24). Such differing compositions engender different availabilities of carbon and nutrients, which, in turn may impact the accompanying bacterial community. Indeed, the phytoplankton host species may drive bacterial community dynamics (24-27) and shape the accompanying bacterial community (15, 28-30). The magnitude of interference of the phytoplankton host species on DOM utilization of the accompanying bacterial community, however, is still subject of debate. Earlier experiments with DOMp addition derived from cyanobacteria and diatoms revealed overall highly similar transcriptomic profiles of bacteria (31), and the two contrasting phytoplankton groups sometimes trigger the same bacterial communities in bloom events (28). On the other hand, only one out of many *Prochlorococcus* strains could meet the unique glycine requirements of SAR11 (32), and in the before mentioned studies, some bacteria were specialized in either cyanobacteria or diatom derived DOM (31), and responsive bacteria differed between other bloom events of diatoms or cyanobacteria (28). A second aspect that needs to be considered if analyzing bacterial utilization of DOMp is the temporal transformation of compounds and the concomitant succession of specific bacterial phyla after DOM pulses (28, 33-35). The DOMp composition changes with proceeding time from highly bioavailable, mostly low-molecular-weight (LMW) to less bioavailable, mostly high-molecular-weight (HMW) compounds, caused by the preferential utilization of labile LMW compounds by bacteria (36). Analogue to the phytoplankton source specificity of specific bacteria, different bacteria phyla are specialized in the utilization of different DOMp compounds. Typically, members of the Flavobacteria are the principle consumers of HMW compounds, whereas the genetic endowment of the Alphaproteobacterial Roseobacter clade reflect their preferences for LMW compounds (28, 31, 33, 34).

However, also the same bacterial phyla may adapt to the transformation of DOMp compounds by changing expression patterns of transporters (37), and various HMW compounds of the same molecular weight may reveal tremendous differences in their accessibility due to compositional and structural differences (38).

In a previous set of experiments, we showed that cyanobacteria DOM provide carbon, nitrogen and phosphorus to accompanying heterotrophic bacteria, and that the uptake of nitrogen and phosphorus bound to DOMp is preferred over inorganic forms (7). Here, we further explore how the interplay of DOM donor species and time after DOMp pulses impact the accompanying bacterial community dynamics. Therefore, we addressed the following questions: (i) do diatom and cyanobacterium DOM serve as P and C source in equal measure? (ii) to which magnitude does the phytoplankton donor species impact the accompanying bacterial community? (iii) how do bacterial communities change over time following DOMp pulses? (iv) which are the primary active bacteria in DOMp utilization from both phytoplankton donor species, and (v) does the specificity of heterotrophic bacteria to the phytoplankton donor species change with DOMp transformation over time?

In order to begin answering these questions, we performed DOMp addition experiments with two globally abundant phytoplankton species - the diatom *Skeletonema marinoi* and the cyanobacterium *Prochlorococcus* MIT9312 - over a time-course of 72 h. Diatoms and cyanobacteria are the two primary sources of marine organic carbon (39), where *Prochlorococcus* contributes up to 8.5% of oceanic photosynthesis and 50% of surface open-ocean chlorophyll (40), whereas diatoms execute up to 40% of primary production in coastal areas (41).

## RESULTS

### Nutrient concentrations

To test for possible nutrient limitations during the course of the experiment, we measured PO_4_, NO_x_ and NH_4_ concentrations. PO_4_ concentrations were ca. 0.1-0.2 µM in the control and the diatom DOM treatments, whereas ca. 1 µM PO_4_ were accidently added with the cyanobacterium DOM. However, neither cyanobacterium DOM treatments with high PO_4_ concentration, nor control or diatom DOM treatments showed pronounced changes in PO_4_ concentrations over the course of the experiment (Supplemental Fig. 1). NO_x_ concentrations at the start of the experiment were ∼ 2.5 µM and remained constant throughout the entire experiment in control and cyanobacterium DOM treatment, but decreased after 48 and 72 h in the diatom DOM treatment (Supplemental Fig. 1), whereas NH_4_ concentrations ranged between 1 and 15 µM but did not reveal obvious trends over time. Nevertheless, control samples showed slightly higher NH_4_ concentrations compared to the DOMp (diatom and cyanobacterium) treatments, which were, however, significantly higher only after 24 h of exposure (Supplemental Fig. 1).

### Bacterial cell numbers, Bacterial Production and Alkaline Phosphate Activity

Initial bacterial cell numbers of winterly, coastal Mediterranean waters were ca. 2.25 million cells ml^-1^, and elevated after 24 h of exposure in all treatments, but levelled off to the original value after 48 and 72 h in both DOMp treatments, and decreased to ca. 1.8 million cells ml^-1^ in the control treatment (Fig. 1A).

On the other hand, bulk measurements of bacterial production (BP) increased in all but the control treatments over the course of the experiment, yet differed significantly between the different treatments at all time-points (Fig. 1B). After 24 h, BP in the diatom DOM treatment was higher than BP in the controls and the cyanobacterium DOM treatments, whereas after 48 and 72 h both phytoplankton DOM treatments yielded similar BP which were significantly higher compared to the control treatment (Fig. 1B). To test for phosphorus starvation in the individual treatments, we measured the production of alkaline phosphatase activity (APA). Bulk measurements of APA yielded lower activity in the DOMp treatments compared to the controls after 48 and 72 h, without any statistical differences between both DOMp donors (Fig. 1C).

**Fig. 1.**
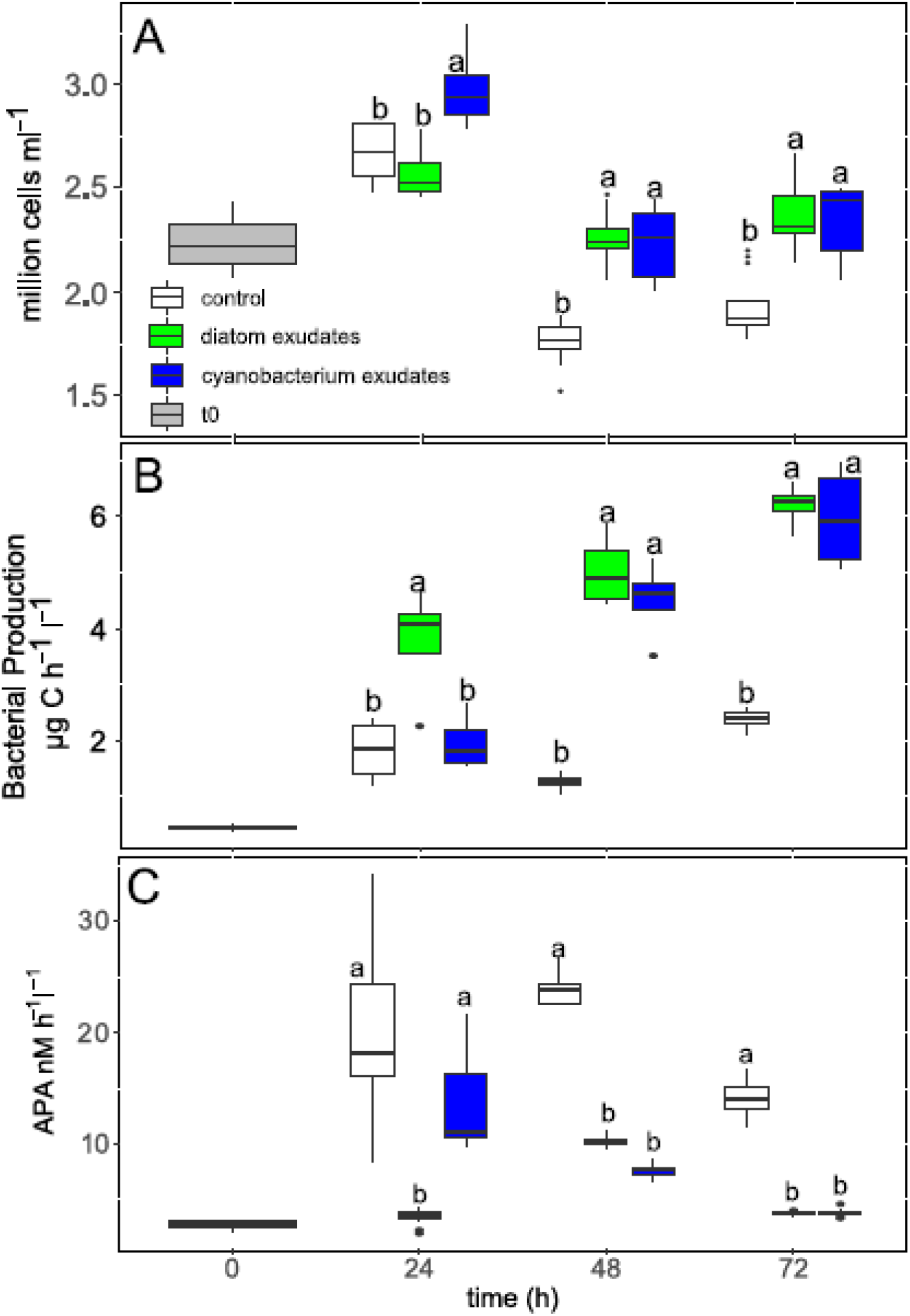
(A): Bacterial cell numbers, (B) Bacterial Production, and (C) Alkaline Phosphatase Activity (APA) for different time-points and treatments. The letters in the panels represent the outcomes of Tukey post-hoc tests between the treatments for each individual time-point (see Supplemental statistics for further information).

However, after 24 h, diatom DOM revealed ca. 4-times lower APA concentrations compared to the cyanobacterium DOM (Fig. 1C).

### Bacterial community responses to DOMp additions

In order to evaluate consequences of different DOMp donors and the succession of DOMp on the richness of the active bacterial communities we calculated Shannon diversities for the individual treatments and time-points. In all samples, the diversity remained in a narrow margin with values ranging from ca. 5.4 to 6.2 (Fig. 2).

**Figure 2:**
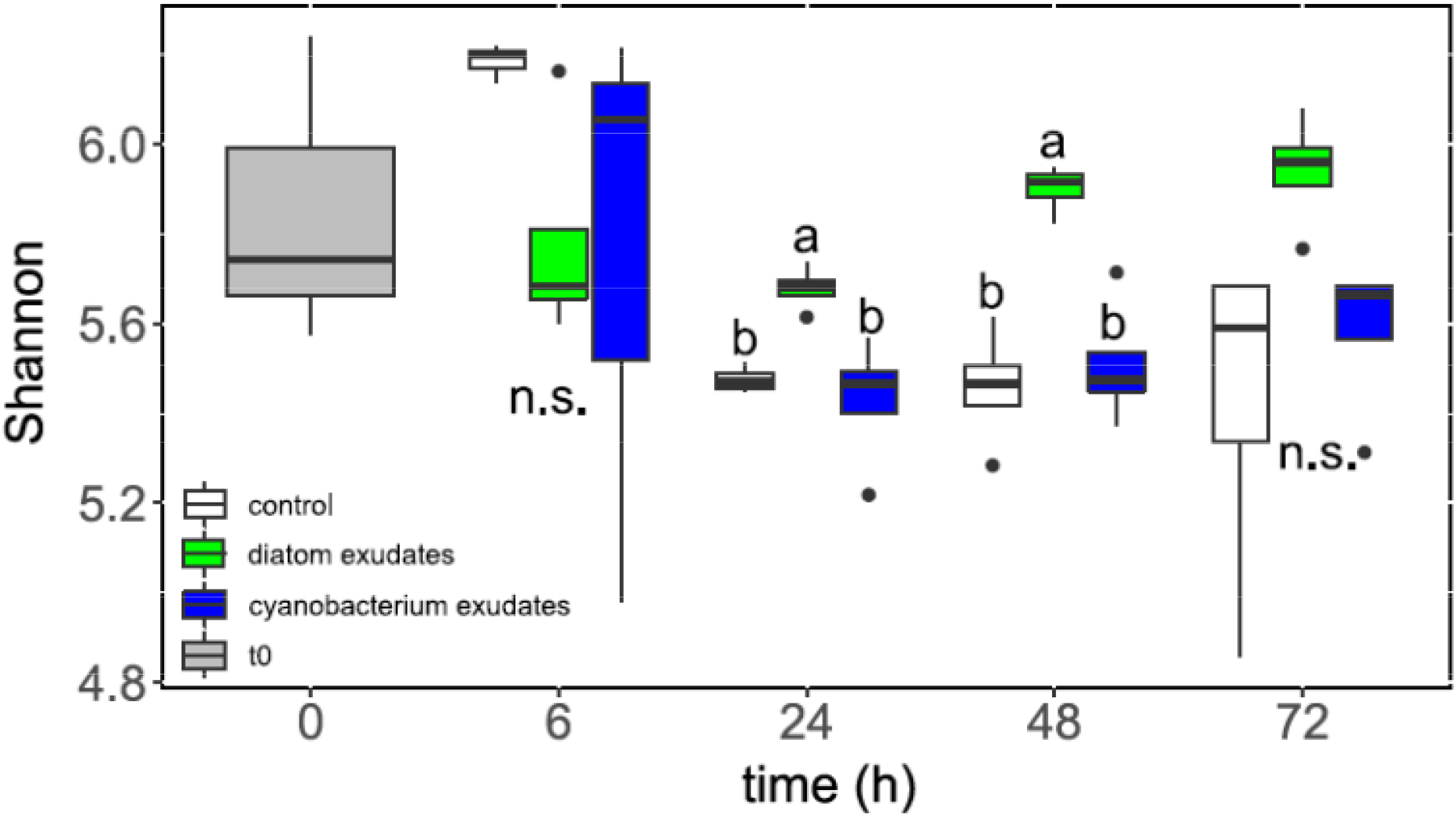
Shannon diversities for the different time-points and treatments. The letters in the panels represent the outcomes of Tukey post-hoc tests between the treatments for each individual time-point (see Supplemental table 2 for further information).

Nevertheless, after 24 and 48 h, samples in the diatom DOM treatments revealed significantly higher diversities compared to cyanobacterium DOM and control samples at this time point (Fig. 2). The increased diversity in diatom samples was consistent also after 72 h of incubation, but not anymore in a significant way (p=0.055, Supplemental table 2). Next, we asked for shared (generalist) and unique (specialist) ASVs between treatments at different time-points using Venn analyses. If shared ASVs between all treatments (control, diatom DOM, cyanobacterium DOM) were compared with each other, no trend over time was obvious (13%, 22%, 20% and 20% after 6, 24, 48, and 72 h of incubation, respectively), whereas ASVs that were only shared between both DOMp treatments (i.e. DOMp generalists) showed a slight increase over time with shares of 2.3%, 3.5%, 4.3% and 4.4% for 6 h, 24 h, 48 h, and 72 h of incubation, respectively (Supplemental Fig. 2). On the other hand, at the different time points (6, 14, 48, 72 h) exclusive (i.e. specialist) ASVs in cyanobacterium (22%, 20%, 20%, and 19%) and diatom (18%, 23%, 26%, and 25%, respectively) DOMp revealed slightly decreasing (cyanobacterium) and variable (diatom) trends over time (Supplemental Fig. 2).

We then explored the impact of the different treatments and progressing time on the active bacterial communities with non-metric multidimensional scaling (NMDS) and Bray-Curtis dissimilarities. Different treatments (t0, control, diatom DOM, cyanobacterium DOM Fig. 3A) showed significantly different bacterial communities (ANOSIM, R=0.3, p=0.001), and this difference remained if only bacterial communities between both DOMp types were compared (ANOSIM, R=0.22, p=0.001, Fig. 3A). However, if treatments were pooled together and samples grouped into different incubation times (0 h, 6 h, 24 h, 48 h, and 72 h), significant differences in communities between the different time-points could be detected (ANOSIM, R=0.4, p=0.001, Fig. 3A).

**Figure 3:**
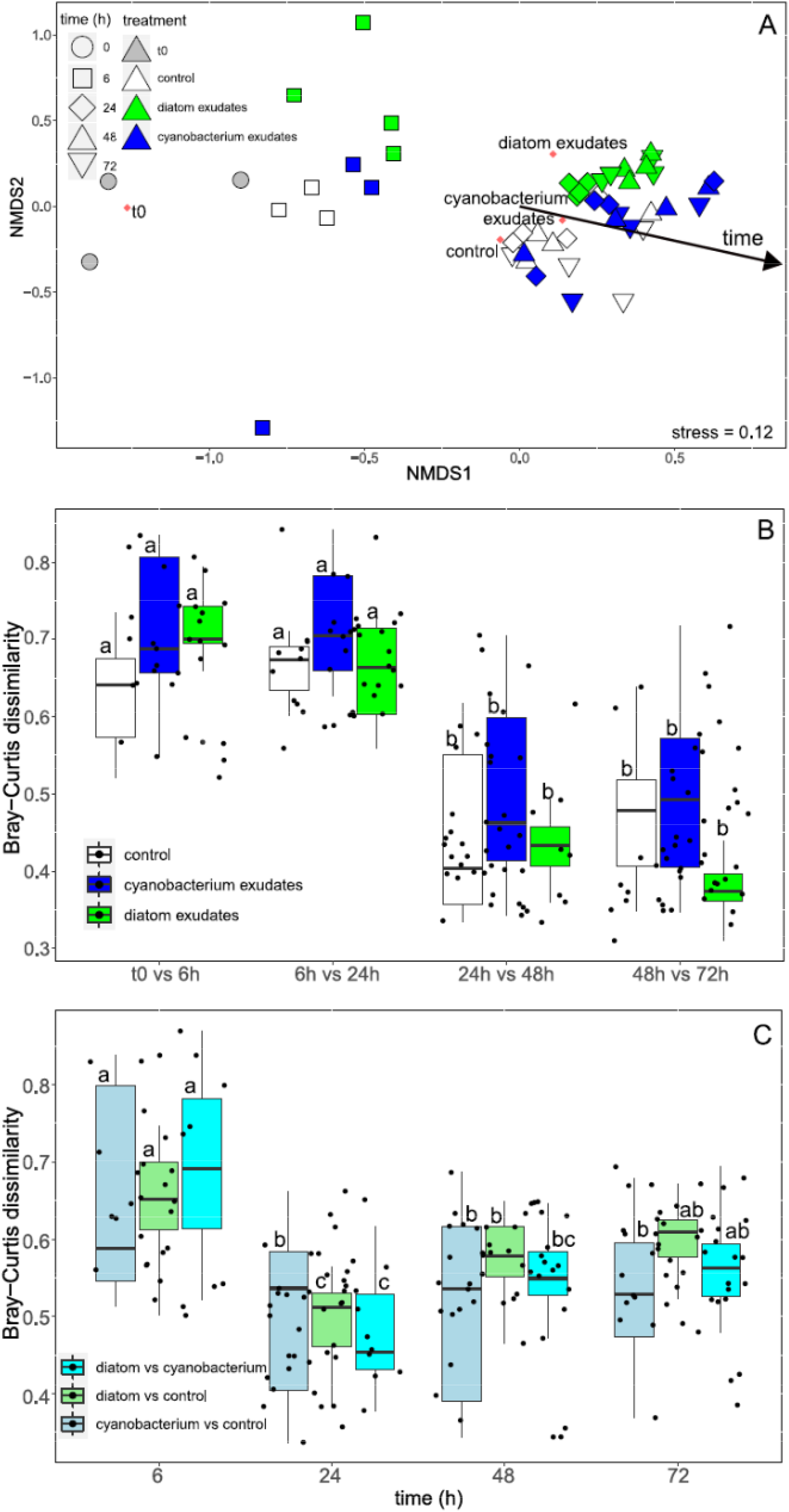
Effects of different treatments and proceeding time on bacterial communities. (A) Non-metric multidimensional scaling (NMDS) for the different treatments and time-points combined with EnvFit analyses. The red squares indicate the calculated centres of the respective treatments, the arrow indicates the direction of proceeding time, (B) Bray-Curtis dissimilarities of bacterial communities between different time-points in the same treatment, and (C) Bray-Curtis dissimilarities of bacterial communities between different treatments at the same time-point. (B, C) Letters on top of the box-plots refer to the outcomes of Tukey post-hoc tests for differences over time in the same treatment (B) and for differences of time-dependent differences between different treatments (C). Further information and additional tests (different treatments, same time-point comparisons (B), and different treatment comparisons at the same time-point (C)) are provided in Supplemental Table 2.

In all treatments, time-dependent changes in the active bacterial community were significantly higher in the first 24 h of exposure (Bray-Curtis dissimilarities between t0 and 6 h as well as between 6 h and 24 h), compared to dissimilarities between later time-points (Figure 3B). Analogue to comparisons between different time-points in the same treatments, community differences between different treatments at the same time point were most pronounced already after 6 h, and the overall highest difference was obtained between diatom and cyanobacterium DOM treatments (Fig. 3C). The differences between bacterial communities in diatom and cyanobacterium DOM treatments, however, diminished after 24 h, but successively increased again after 48 and 72 h (Fig. 3C).

General patterns revealing bacterial community differences between different treatments and incubation times were confirmed by abundance distributions of the overall 100 most abundant ASVs (defined as the sums of the subsampled reads for all samples, Fig. 4). A dendrogram of the ASVs yielded three major clusters, which were largely driven by the taxonomy, with *Sachharospirillaceae, Spongiispira*, SAR11 and KI89a ASVs in cluster I, *Ascidiaceihabitans, Glaciecola, and Aureimarina* ASVs dominating cluster II, and a diverse consortium of ASVs (including *Synechococcus*, Rhodobacteraceae, and others) in cluster III (Fig. 4). However, if the 100 most abundant ASVs were analysed for the dominant ASVs in the different treatments (t0, control, cyanobacterium DOM, diatom DOM), t0 samples were dominated by ASVs from SAR11 and *Synechococcus*, whereas 11 ASVs of *Saccharospirillaceae*, one *Spongiispira* ASV and one *Glaciecola* ASV dominated the control treatments (Fig. 4, SupplementalFig. 3). On the other hand, the cyanobacterium DOM treatments were exclusively dominated by ASVs of *Glaciecola*, whereas two Flavobacteriaceae, one *Ascidiaceihabitans* ASV, one SAR11 ASV, and four *Glaciecola* ASVs were dominating the diatom DOM samples (Fig. 4, Supplemental Fig. 3A). Additional tests for DOMp generalists, i.e. ASVs that dominate DOMp samples independent of the donor species (assigned treatments: all samples containing DOMp, t0, control), yielded 21 *Glaciecola* and two *Flavobacteriaceae* ASVs (Supplemental Fig. 3B). If different incubation times were analysed for their dominant ASVs, t0 samples were defined by a broad array of bacterial genera and ASVs, with the phototrophic *Synechococcus*, and heterotrophic SAR11, OM43 clade, *Ascidiaceihabitans*, AEGEAN-169, NS2b, NS4, NS5, and NS9 marine group contributing several ASVs. Six-hour incubation samples were especially dominated by ASVs of the Rhodobacteraceae, 24 h incubations solely by *Glaciecola* ASVs, 48 h incubations by seven ASVs of *Glaciecola* and one ASV from *Pseudoalteromonas*, and the 72 h incubations by 11 ASVs of Saccharospirillaceae and one *Aureimarina* ASV (Fig. 4, Supplemental Fig. 3C).

**Figure 4:**
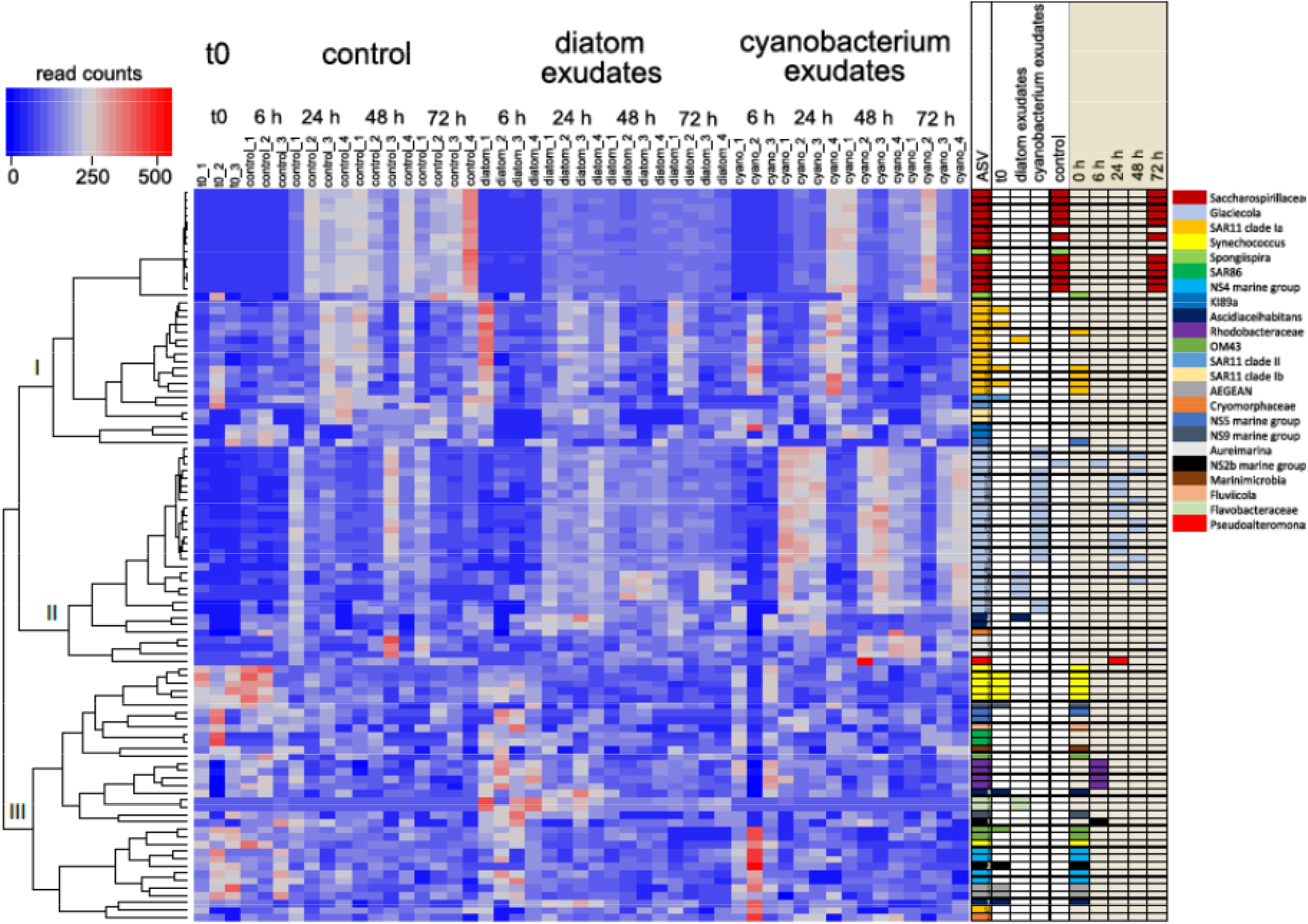
Heatmap of the overall 100 most abundant ASVs for the different treatments and time-points. The dendrogram on the left side of the heatmap is based on hierarchical clustering of ASVs. The table on the right side of the heatmap provides ASV taxonomy (first column with colour code), and dominant ASVs for the different treatments (t0, diatom DOM, cyanobacterium DOM, control, backed with light-grey) as well as different time-points (0 h, 6 h, 24 h, 48 h, 72 h, different treatments pooled, backed with brown-grey, Supplemental Fig. 3A, C). Dominant ASVs of the treatments/time-points are indicated by filled boxes in the colour of the respective taxonomy. ASVs of the same genus are defined by numbers in the taxonomic colour code.

## DISCUSSION

In this study, we confirmed the bacterial utilization of DOMp derived from two phytoplankton primary sources of marine organic matter, i.e. diatoms and cyanobacteria (39) in coastal waters of the Eastern Mediterranean in winter. We found similar bacterial responses to diatom and cyanobacteria DOMp additions in terms of bacterial cell numbers, bacterial production (BP) and alkaline phosphatase activity (APA). Nevertheless, faster bacterial responses in BP and APA following the addition of diatom DOM suggest better accessibility of carbon and phosphorus from these DOMp compared to cyanobacterium DOM. Furthermore, via 16S rRNA sequencing we could demonstrate that different, DOMp-donor specific bacterial ASVs are active in DOMp utilization. However, the most pronounced differences between active bacterial communities of both DOMp types were obtained shortly after additions (6 h), indicating an important role of donor-specific DOMp compounds in the highly bioavailable DOMp fraction. We also observed bacterial community successions over time, which we hypothesize are due to a shift from ASVs effective in utilization of highly bioavailable, donor-specific DOMp compounds shortly after the DOMp pulse, towards ASVs specialized on less bioavailable compounds at the later time points. In summary, we conclude that bacterial communities experiencing phytoplankton DOMp pulses are relevantly defined by an interplay between the original DOMp composition and temporal succession in substrate quality following the DOMp pulse.

### Effects of DOMp additions on Bacterial Cell Numbers, Bacterial Production and Alkaline Phosphatase Activity

We confirmed bacterial utilization of phytoplankton DOMp as carbon source with measurements of bacterial cell numbers and production (BP) (Fig. 1), and found indications for phosphorus utilization due to lowered alkaline phosphatase activity (APA) in DOMp treatments (Fig. 1). Absolute cell-numbers appear to be surprisingly high for coastal Mediterranean Waters, with ∼2.2 million cells ml^-1^ at t0 (Fig. 1A). Yet, these values correspond well to our previous study in winter (also ∼2.2 million cells ml^-1^ at t0), F. Eigemann et al. (7)), as well as with a time series in the Eastern Mediterranean (∼0.7 – 1.4 million cells ml^-1^ in January/February, O. Raveh et al. (42)), both sampled from the same station. Likewise, increased BP in both DOMp treatments corroborated bacterial utilization of phytoplankton derived DOC, with well corresponding values to O. Raveh et al. (42) (∼0.8 – 1.6 µg C h^-1^ l^-1^ in January/February vs ∼0.5 µg C h^-1^ l^-1^ at t0 in the present study), and our previous experiment (t0 ∼0.13 fg C cell^-1^ h^-1^, F. Eigemann et al. (7) vs ∼0.2 fg C cell^-1^ h^-1^, this study). Significantly lower APA in diatom DOM treatments compared to controls despite of identical inorganic P concentrations (Fig. 1C, Supplemental Fig. 1), suggests that diatom DOM wwas used as an organic P source. This is in accordance with findings that bacterial communities in the Mediterranean Sea compensate for low inorganic nutrient levels with organic forms (43, 44), and phytoplankton DOMp represent a substantial source of it (45). On the other hand, we added approximately 1 µM PO_4_ with the cyanobacterium DOM (Supplemental Fig 1), which makes a definitive statement on P utilization derived from cyanobacterium DOM difficult. However, in our preceding study we showed a preferentially utilization of organic nutrient forms provided from cyanobacterium DOM compared to inorganic forms (7). In the present study, we may have experienced a similar organic P utilization, which is suggested by identical outcomes for t0 and control APA in both studies. The even higher APA in the cyanobacterial DOM treatments despite of our accidental PO_4_ addition (Fig. 1) in parallel to constant inorganic nutrient levels with contemporaneous decreasing APA over time (Fig. 1, Supplemental Fig. 1) supports this notion.

### Faster accessibility of diatom DOM and donor specific effects on bacterial diversity

DOMp differs in its composition between different phytoplankton groups (16, 21), as well as between species in the same group (13), which has consequences on their accessibility to the bacterial community (23, 24, 39). After 24 h of exposure, bacteria exposed to diatom DOM yielded approximately 2-times higher BP and 2.5-times lower APA compared to bacteria exposed to the cyanobacterium DOM (Fig. 1). This pattern was consistent for bulk as well as per cell measurements, and thus not attributed to the higher cell-numbers in cyanobacterium DOM treatments after 24 h of exposure (Supplemental Fig. 4). Both results seem surprising at a glance, considering that we added less carbon with the diatom than cyanobacterium DOM (14 vs 25 µM, see Experimental procedures), and also added inorganic PO_4_ with the cyanobacterium DOM (see above). Indeed, our results suggest a faster and/or higher accessibility of diatom DOM for bacterial biomass synthesis and phosphorus utilization, which is in accordance with the outcomes by B. Kieft et al. (39), who found a higher ^13^C enrichment in bacteria exposed to^13^C-labelled diatom than cyanobacterium DOM after 15 h, and a mass spectrometric analysis of both DOMp types, where diatom DOM owned a higher fraction of P (23).

Higher Shannon diversities throughout the experiment and a higher diversity of genera (and not ASVs) in the 50 most abundant ASVs in the diatom DOM samples (Supplemental table 3), furthermore suggest that diatom DOM can be used by a larger fraction of the bacterial community compared to the cyanobacterium DOM (Fig. 2). This outcome may be attributed to the production of easy accessible DOM such as the polysaccharide laminarin (46), and more evenly distributed individual sugars and amino acids in diatom DOM (which infer a higher diversity of polymers, and thus can be used by a broader share of bacteria) compared to cyanobacteria DOM (23), which also contain lipo-polysaccharides (47), as well as fucose containing polysaccharides (23). The latter requires sophisticated degradation pathways.

Nevertheless, cyanobacterium and diatom DOM treatments experienced different DOC und P concentrations, which might have impacted bacterial responses. Higher APA in higher PO_4_, and lower BP in higher DOC environments, however, indicate a DOM quality (i.e. source) and not quantity driven response. Further, higher DOC concentrations were, independent of the phytoplankton donor, utilized by a larger fraction of bacteria (48), suggesting that the higher bacterial diversity in our diatom vs. cyanobacterium DOM treatments indeed derives from DOMp quality and not quantity.

### DOMp donor specific ASVs

We found multiple different active ASVs of Alphaproteobacteria, Gammaproteobacteria, Bacteroidetes, Verrucomicrobia, Firmicutes, Marinimicrobia, Latescibacteriota, and Actinobacteria in samples of both phytoplankton donors as abundant members (Supplemental table 3). The vast majority of abundant ASVs in DOMp enriched samples, however, were assigned to Gammaproteobacteria, the Bacteroidetes family Flavobacteriaceae, or the Rhodeobacterales clade of Alphaproteobacteria (Supplemental table 3). These three bacterial groups are primary utilizers of DOMp and are, independent from the phytoplankton donor species and time after DOMp pulses, well known as dominant members in associations with phytoplankton, and/or bacterial communities after phytoplankton DOMp additions (27, 28, 32-34, 39, 49-51).

At the ASV level, however, we determined 17 different *Glaciecola* ASVs in cyanobacteria and four different *Glaciecola* ASVs in diatom DOM samples dominating the respective treatments (Supplemental Fig. 3). The assignment of only *Glaciecola* (Gammaproteobacteria, Alteromonadaceae) ASVs to cyanobacterium DOM samples is very similar to the outcomes in our previous study (7), and also corroborated by studies of H. Sarmento and J. M. Gasol (51) and S. M. Kearney et al. (52), who found preferential uptake of Prochlorococcus DOM by Alteromonadaceae, and specific associations of Alteromonadaceae to Piccocyanobacteria, respectively. However, *Glaciecola* ASVs were assigned to both DOMp types (Supplemental Fig. 3A, 3C), and obtained high relative abundances in conditions dominated by different phytoplankton species (53), and thus *Glaciecola* might be considered as a generalist phytoplankton associated bacterium (or DOMp generalist, Supplemental Fig. 3B). Besides of four *Glaciecola* ASVs, diatom DOM samples were dominated by two Flavobacteriaceae, and each one SAR11 and *Ascidiaceihabitans* ASVs (Supplemental Fig. 3A). These assignments are supported by numerous studies that found general associations of Proteobacteria and Bacteroidetes towards diatoms (reviewed by S. Amin (54)). However, despite of several specific phytoplankton-bacteria associations that were found in our and other studies, one should be careful to draw general conclusions. First, we indeed defined several ASVs as phytoplankton donor specific, but one should keep in mind that also other ASVs were abundant in either diatom or cyanobacteria samples, whose non-consideration as donor specific might be a result of statistical cut-offs, which may not reflect soft boarders in natural systems. Second, divergent outcomes were obtained in similar studies. As examples, we predominantly assigned *Glaciecola* to cyanobacteria DOM, whereas M. Landa et al. (23) found it specific to diatom DOM (and did not find it in cyanobacteria DOM treatments). Further, Rhodobacterales were specifically associated with picocyanobacterial (39, 52), and Prochlorococcus supported the growth of SAR11 in co-culture experiments (32), whereas we assigned *Ascidiaceihabitans* (Rhodobacterales) and SAR11 only to diatom treatments. Thus, general assignments of specific phytoplankton-bacteria associations are challenging, which might be attributed to differences in abiotic parameters (which determine the majority of bacterial associations with phytoplankton (55)) as well as bacterial source communities (which greatly impact associations with phytoplankton (56)) between studies.

### Diminishing impact of the DOMp donor on bacterial communities over time

We observed the most pronounced bacterial community changes between 0 and 6, as well as 6 and 24 h of exposure (Fig. 3B), suggesting a fast operation of treatment specialized bacteria. Nevertheless, successions of the bacterial community occurred throughout the complete incubation period (see Bray-Curtis dissimilarities between 48 and 72 h of exposure, Fig. 3B), and we were able to identify dominating ASVs for each individual time-point (Supplemental Fig. 3C). Bacterial community successions after DOMp pulses follow the exploitation of different DOMp compounds over time, from highly bioavailable compounds shortly after the pulse towards less bioavailable compounds at later time-points (33). The broad consortium of different bacteria dominating t0, changed to four Rhodobacter (Alphaproteobacteria) ASVs and each one *Glaciecola* (Gammaproteobacteria) and NS2b marine group (Flavobacteriaceae) ASVs dominating the 6 h exposure (Supplemental Fig. 3C). The high share of Alphaproteobacteria at this time-point underlines their task as first responders after DOMp pulses, possibly as a result of their high expression of ABC and TRAP transporters that enable the uptake of various LMW compounds (33, 39). After 24 and 48 h, we only assigned Gammaproteobacteria (and only one none *Glaciecola* ASV), and after 72 h eleven Gammaproteobacteria (all Saccharospirillaceae), and only one Flavobacterium as time-point dominating (Supplemental Fig. 3C).

Interestingly, these outcomes exactly reflect the patterns from H. Teeling et al. (33), who also found a succession of peaking Gammaproteobacterial clades, after the primary response of Alphaproteobacteria to a diatom bloom. The secondary peak of Gammaproteobacteria and to a lesser extent Flavobacteria is a recurrent phenomenon after DOMp pulses (34), and can be explained by their genetic endowment that enable the utilization of numerous HMW compounds (33, 57).

The donor specific responses as well as bacterial community differences between both DOMp types were most pronounced shortly after DOMp additions (Figs. 1, 3), suggesting a strong impact of the donor type shortly after DOMp addition. We hypothesize that this is caused by high shares of donor-specific compounds in the highly bioavailable DOMp fractions, whereas less bioavailable compounds may have a more general composition. This assumption is supported by increases in shared ASVs between both DOMp types over time (when highly bioavailable compounds decrease, Supplemental Fig. 2), the decrease of Bray-Curtis dissimilarities from 6 to 24 h of exposure (Fig. 3C), and the different outcomes for BP, APA and cell numbers after 24, but not after 48 and 72 h (Fig. 1). Contradictory, M. Landa et al. (23) suggested that labile compounds have a minor effect on the bacterial community and that pronounced effects of the donor species occur if enzymatic machineries are required. In their study, however, a continuous supply of cyanobacterium as well as diatom DOM was applied, which may have masked a succession from highly to less bioavailable compounds. Furthermore, lower recurrences of phytoplankton species in blooms, compared to heterotrophic bacterial OTUs days to weeks after the blooms between years (34), suggest high similarities in the less bioavailable fraction of phytoplankton DOM derived from different species, and supports our conclusions on DOMp generalists (Supplemental Fig. 3B). Finally, a mechanistic interference study on the same dataset revealed ∼3-times higher recurrences of DOM species compared to phytoplankton species between years in spring blooms (58), which, taking into account that highly bioavailable compounds may be rapidly consumed, further supports our hypothesis.

### Limitations and synthesis

Despite of reconstructing the occurring environmental conditions in our mesocosm experiments, we experienced pronounced bottle effects, illustrated by increased bacterial production in the controls (Fig. 1B), similar temporal developments of bacterial communities in all treatments (including the controls, Fig.3A), and an increase in shared ASVs between all treatments with increasing time (Supplemental Fig. 2). However, first, it is almost impossible to completely omit bottle effects in these types of experiments (e.g., S. Elovaara et al. (59) or C. M. Luria et al. (60)).

Second, despite of obvious changes in control communities from t0 to 6 h of incubation, the highest community differences for these time steps were achieved between both DOMp treatments (Fig. 3C). Furthermore, we found pronounced differences between the phytoplankton donors in APA, BP, and cell numbers after 24 h of exposure (Fig. 1), and were able to define phytoplankton donor-specific ASVs, as well as DOMp generalist ASVs (Supplemental Fig. 3B). Thus, the observed bottle effects did not prohibit the gain of valuable information.

As synthesis, our results and that of similar studies suggest that bacterial diversity, Alkaline Phosphatase activity (APA), Bacterial Production (BP), and cell numbers of phytoplankton accompanying bacterial communities are determined by the phytoplankton donor species (Fig. 5), which may be caused by different DOMp compositions of cyanobacteria and diatoms (20). Additionally, the bacterial community participates in the succession of DOMp from highly bioavailable to less bioavailable (33, 61), with time-point defining ASVs (Supplemental Fig. 3C). However, decreasing differences with increasing time of bacterial communities in the same treatment (Fig. 3B), as well as high community differences between both DOMp treatments after 6 h incubation (Fig. 3C), suggest a particularly high impact of the host shortly after DOMp additions (Fig. 5). After exploitation of species-specific highly bioavailable compounds, the remaining DOMp of different phytoplankton may become more similar, and bacterial communities might be especially impacted by the DOM accessibility (Fig. 5), i.e. the state of DOMp succession from highly bioavailable to less bioavailable compounds.

**Fig. 5:**
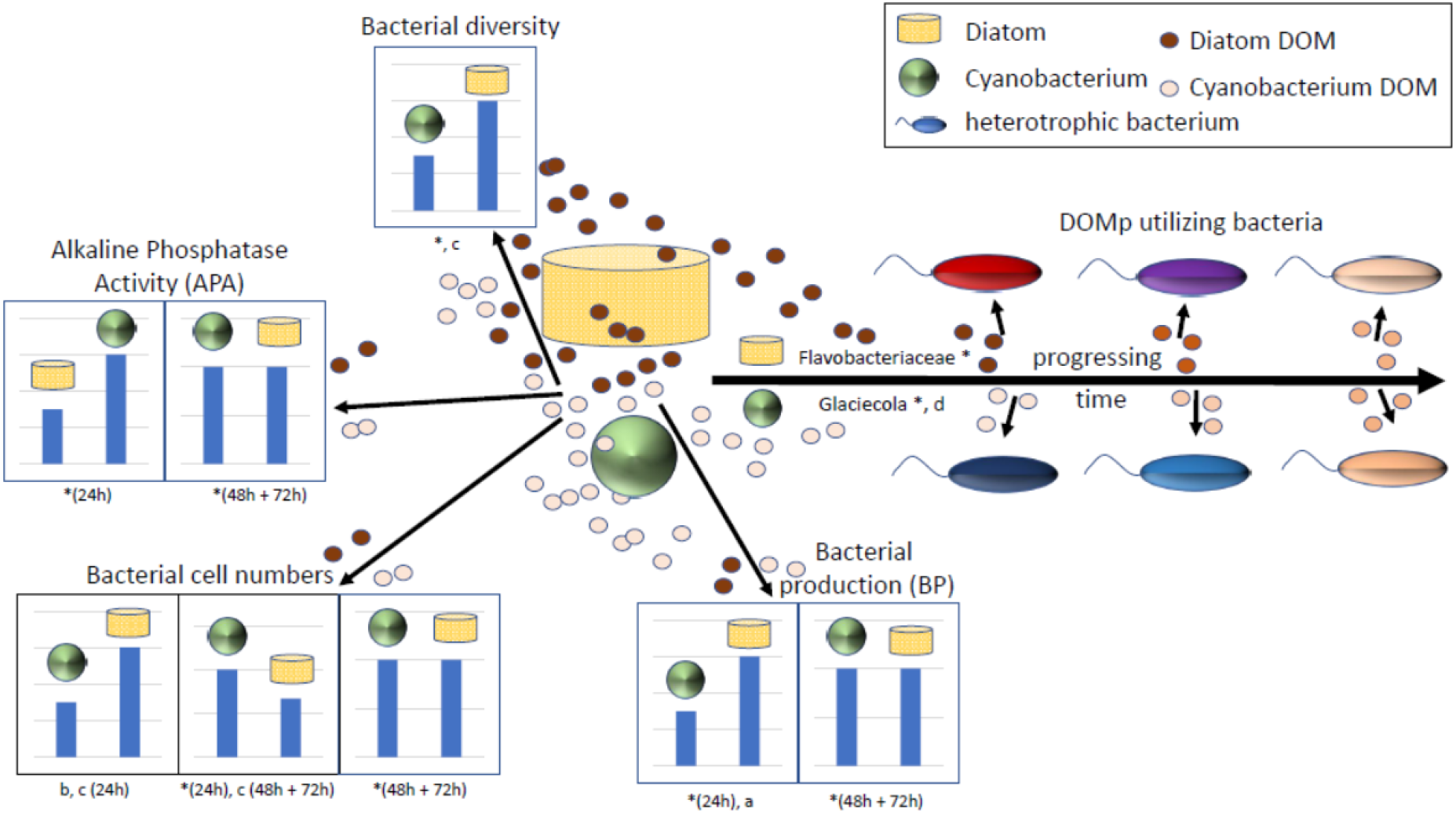
Summary figure of the obtained results (indicated by *), and outcomes of similar studies on bacterial utilization of cyanobacteria and diatom derived DOM (a = B. Kieft et al. (39), b = H. Sarmento and J. M. Gasol (51), c = M. Landa et al. (23), d = F. Eigemann et al. (7)). If different outcomes at different time-points were obtained, a time-specification is given after the reference. Right side: Different colored bacteria indicate taxonomically different bacteria, with higher similarities between colors indicating higher taxonomical similarities. The approximating colors of diatom and cyanobacteria DOM with progressing time indicate the succession of DOMp compounds from highly to less bioavailable, which might become more similar.

## MATERIALS AND METHODS

### Experimental set-up

The experiment was performed in winter from 14. January 2019 until 17. January 2019 at the Israel Oceanographic and Limnological Research center in Haifa, Israel. Shaded 20 l Nalgene bottles were filled with 18 l of 50 µm pre-filtered sea-water (intake pipe at 5 m depth, ca. 50 m off the shore), amended with phytoplankton DOM, and placed in 1 m³ natural seawater flow-through tanks to maintain temperature. We added 25 µM cyanobacterium (*Prochlorococcus* MIT9312) DOC, in order to produce comparable outcomes with similar studies (see F. Eigemann et al. (7)), and 14 µM diatom (*Skeletonema marinoi*) DOC (due to technical issues, several DOMp containing bottles broke at an air-transfer). Each of the three treatments (control, diatom DOM, cyanobacterium DOM) contained four biological replicates, and samples were taken at t0 (nutrients, RNA, APA, BP, cell-numbers), after 1 (only nutrients), 6 (nutrients and RNA), 24, 48 and 72 h (APA, BP, RNA, cell-numbers, nutrients). A graphical overview of the experimental set-up is given as Fig. 6.

**Figure 6:**
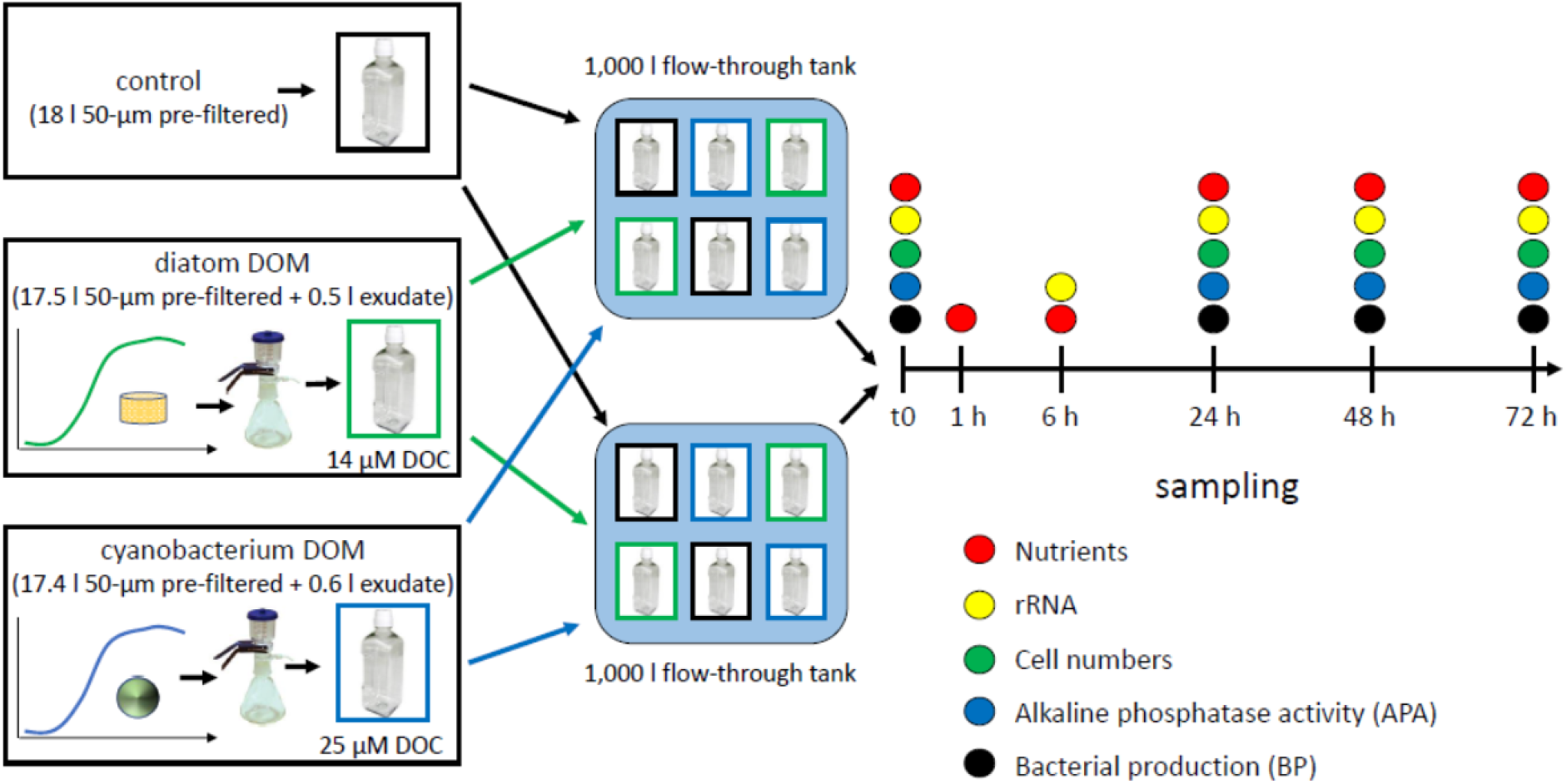
Schematic overview of the experimental set-up. *Prochlorococcus* MIT9312 (cyanobacterium) and *Skeletonema marinoi* (diatom) were grown till the early decline phase, cell-free DOMp harvested, and mixed with pre-filtered natural water. Incubation bottles were placed in darkened flow-through tanks.

### Culture conditions and harvesting of DOMp

*Prochlorococcus* MIT9312 was grown under constant light (20 μ E m^-2^ sec^-1^) at 22°C in Pro99 media where the NH_4_ concentration was reduced from 800 µM to 100 µM, resulting in the cells entering stationary stage due to N starvation (62). *Skeletonema marinoi* was grown in F/2 + Si medium (63) at 45 μ E m^-2^ sec^-1^ at 17°C in a conditioning-cabinet. To obtain cell-free DOMp, batch cultures were harvested at the early decline phase by centrifugation followed by filtration through a 0.22 µm polycarbonate filter. Phytoplankton derived dissolved organic matter (DOMp) from the early decline phase may reflect natural DOMp compositions better than from exponentially growing cultures (12), and includes DOMp from phytoplankton exudation as well as cell lysis. The different DOMp from both phytoplankton were kept at −20°C until utilization.

### Nutrient measurements

Nutrients were measured colorimetrical according to H. P. Hansen and F. Koroleff (64) by means of a Seal Analytical QuAAtro constant flow analyzer. Concentrations of PO_4_, NO_x_ (NO_2_ + NO_3_), and NH_4_ in the different treatments over the course of the experiment are given in Supplemental Fig. 1.

### Cell numbers, Bacterial production, alkaline phosphatase activity, RNA extraction, DNA digestion, cDNA synthesis and sequencing

All analyzes were performed as described in F. Eigemann et al. (7). Brief descriptions of the used techniques and protocols are given below.

### Cell numbers

Duplicate samples were fixed with cytometry-grade glutaraldehyde (0.125% final concentration), flash-frozen with liquid nitrogen and stored at −80 °C until analyses. For measurements, samples were thawed in the dark at room temperature, stained with SYBR Green I (Molecular Probes/ ThermoFisher) for 10 min at room temperature, vortexed, and run on a BD FACSCanto™ II Flow Cytometry Analyzer Systems (BD 146 Biosciences).

### Bacterial production and alkaline phosphatase activity (APA)

Bacterial production was measured using the [4,5-^3^H]-leucine incorporation method (65). Samples were amended with 100 nmol of leucine L^-1^ (Perkin Elmer, specific activity 156 Ci mmol^-1^) and incubated for 4 h in the dark under ambient surface sea water temperature (∼19 °C). Incubations were stopped by the addition of 100 µL of ice-cold 100% trichloroacetic acid (TCA). APA was determined by the 4-methylumbeliferyl phosphate (MUF-P: Sigma M8168) method according to (66).

Substrate was added to a final concentration of 50 µM and incubated in the dark at ambient temperature for 4 h (same as BP).

### RNA extraction, DNA digestion, cDNA synthesis

In order to analyze active bacteria of the bacterial community, rRNA was extracted and transcribed into cDNA. Approximately 1.5 l of each incubation bottle was filtered directly onto 0.2 µm filters, and extracted with TRI reagent. Remaining DNA was digested using the Turbo DNA free kit (Invitrogen) using the manufacturer’s instructions, and successful digestions were tested by PCRs. RNA was transcribed into cDNA using MultiScribe reverse transcriptase following the manufacturer’s instructions (Invitrogen).

### Sequencing and Sequence analyses

Sequencing was performed using primers 515F and 926R (67) and an Illumina MiniSeq mid-output flow cell. Forward and backward sequence reads were deposited at the European Nucleotide Archive under the accession number PRJEB44710. All sequences were analyzed using the Dada2 (68) pipeline and the software packages R (70, 71) and RStudio (72). Briefly, forward and backward primers were trimmed, forward and backward reads truncated after quality inspections to 280 and 210 bases, respectively, and after merging of forward and backward sequences, a consensus length only between 404 and 417 bases was accepted. For taxonomic assignment, Silva database version 138 (73) was used. All chloroplasts, mitochondria, archaea, eukaryotes and Amplicon sequence variants (ASVs) without any taxonomic affiliation were discarded from downstream analyses. The complete ASV table with absolute, subsampled read counts, and meta data for all samples is accessible as Supplemental Table 1. After inspections of rarefaction curves, samples with less than 10,154 reads were discarded from further analyses (t0, replicate four (41 reads), and cyanobacterium DOM, replicate four, 6 h incubation (1,955 reads)), and all remaining samples subsampled to 10,154 reads. One additional sample was discarded from downstream analyses, because of a sandy RNA filter appearance, and NMDS clustering far away from all other samples (control, replicate four, 6 h incubation). Thus, from originally 52 samples, 49 were used for elaborated analyses.

### Statistics

We tested for differences between the treatments at individual timepoints with Shannon indices, cell numbers, Alkaline Phosphatase Activity (APA), and bacterial production. Variance homogeneities between the different treatments for each individual time-point were tested with Levene tests with the R package car (74). If homogeneities were given, ANOVAs with subsequent Tukey post-hoc tests were calculated. If no homogeneities of variances were given, Kruskal-Wallis tests with subsequent Tukey-Nemenyi post-hoc tests were executed with the R package PMCMR (75). To test for unique and shared ASVs between treatments, we merged biological replicates as follows: All subsampled, absolute read counts >0 were set to 1, and subsequently the mean abundance of the biological replicates was calculated for each treatment. An ASV was set as present if the calculated mean for the treatment was ≥ 0.5. With these matrices, Venn counts were calculated and Venn diagrams generated with the limma package (76). Changes in bacterial communities were examined with Shannon diversities, non-metric-multidimensional-scaling (NMDS), Bray-Curtis dissimilarities between treatments and time-points, and heatmaps of the overall 100 most abundant ASVs. Differences of Bray-Curtis dissimilarities between samples were tested with ANOVAs (homogeneities of variances given) or Kruskal-Wallis tests (homogeneities of variances not given), with subsequent Tukey post-hoc or Tukey-Nemenyi post-hoc tests, respectively (see above). NMDS was executed with Bray-Curtis distances in the vegan package (Oksanen et al. 2019) with subsampled absolute ASV reads. To test for differences between bacterial communities, analyses of similarities (ANOSIM) as well as envfit (environmental fits on ordinations) analyses were conducted using the vegan package. Heatmaps for the overall 100 most abundant, subsampled ASVs were executed using the heatmap 3 package (70, 77). To test if specific ASVs of these 100 most abundant ASVs are associated with specific time-points or treatments (control, diatom DOM, cyanobacterium DOM), LDA effect size (LEfSe) analyses (78) were executed with the online tool https://huttenhower.sph.harvard.edu/galaxy. Time-points and treatments were assigned as class, and samples as subjects. Then, a “one-against-all” strategy was applied with normalization and the LDA effect size threshold set to 2.5. All analyses, except denoted otherwise, were executed with R (71) and RStudio (72). Statistical outcomes are summarized in Supplemental Table 2. All graphics, except denoted otherwise, were executed with the ggplot2 package (79), and refined with the freeware Inkscape (https://inkscape.org).

## ACKNOWLEDGEMENT

We thank Stefan Green (DNA Services Facility at the University of Illinois at Chicago) for the amplicon sequencing, and Christian Burmeister for nutrient analyses. We also thank Tom Reich, Dalit Roth-Rosenberg, Tal Luzzatto-Knaan, Noam Nago, and Natalia Belkin for excellent help with the experiment. This work was supported by the Human Frontier Science Program (HFSP) through the grant number RGB 0020/2016 (DS, MV and HPG), by the National Science Foundation - United States-Israel Binational Science Foundation Program in Oceanography (grant number 1635070/2016532 to DS) and by the Israel Ministry of Science and Technology (grant number 3-17404 to DS).

---

**Supplemental Figure 1:**
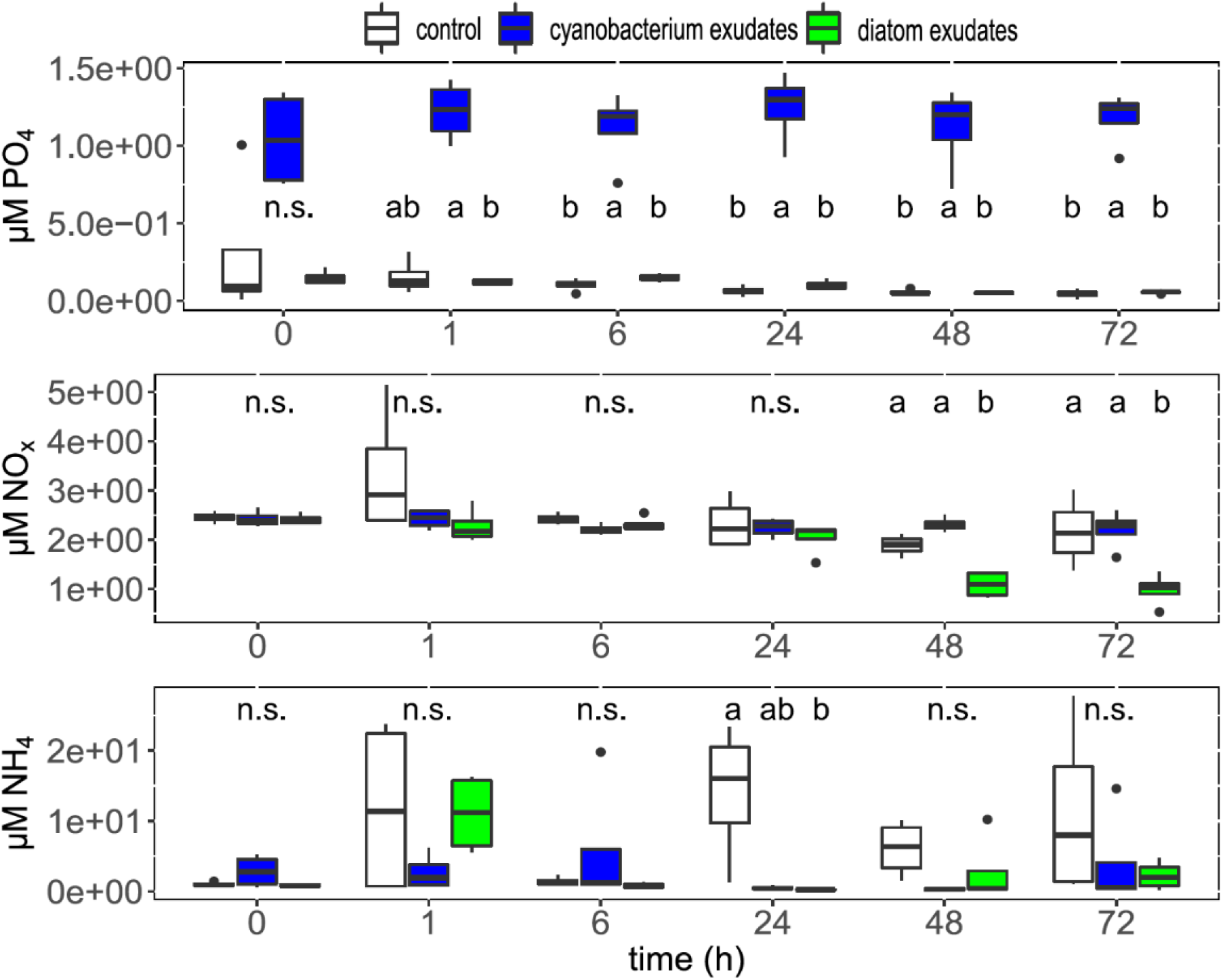
Nutrient concentrations for the different treatments over time. The letters in the panels represent the outcomes of Tukey post-hoc tests between the treatments for each individual time-point (for further information see Supplemental Table 2). If values were below the detection limit, they were set to half of the detection limit for graphical purposes.

**Supplemental Figure 2:**
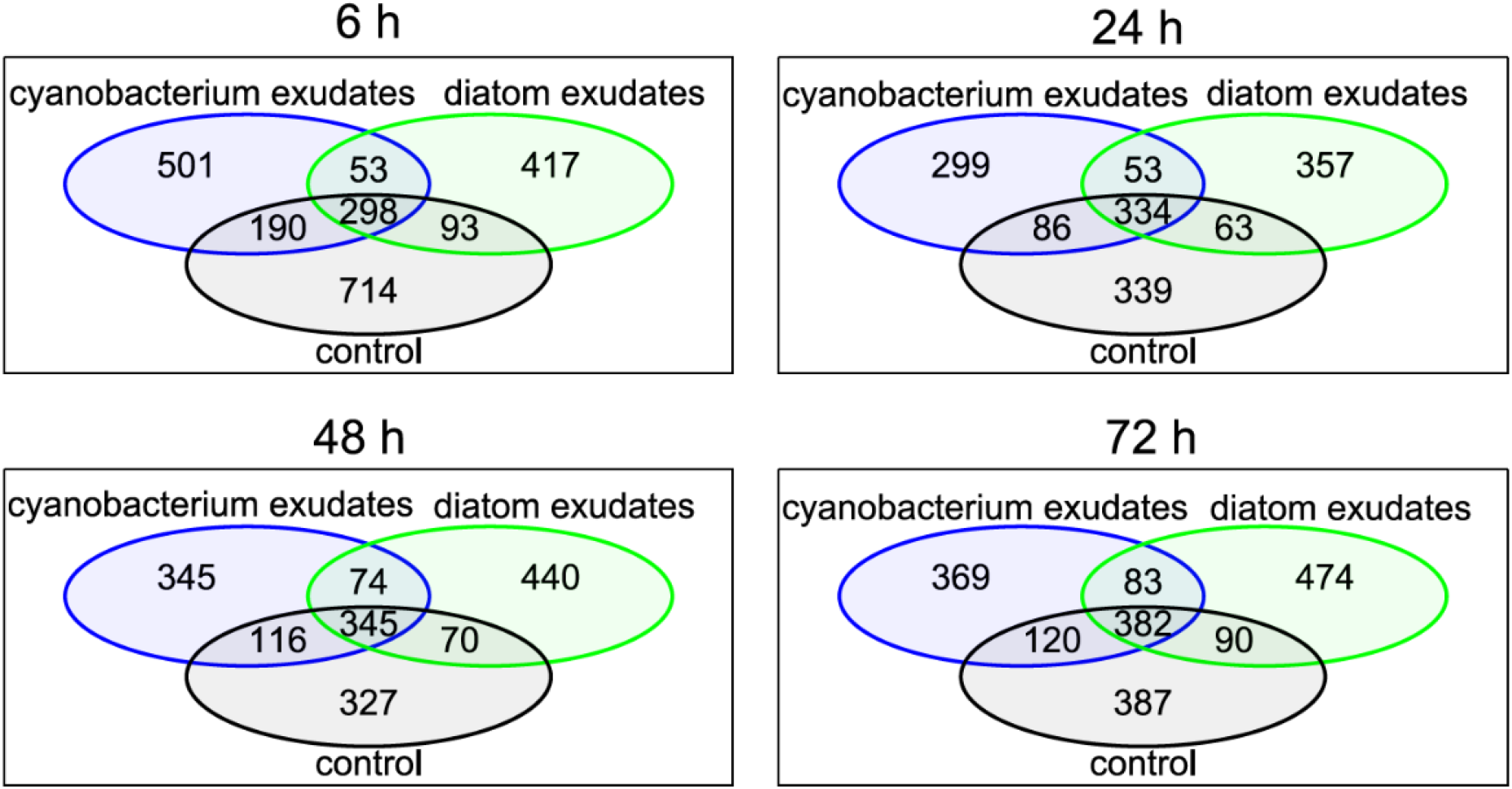
Venn diagrams of shared and unique ASVs for the different treatments and time-points calculated as means of the replicates (see Experimental procedure for details).

**Suppementary Figure 3A:**
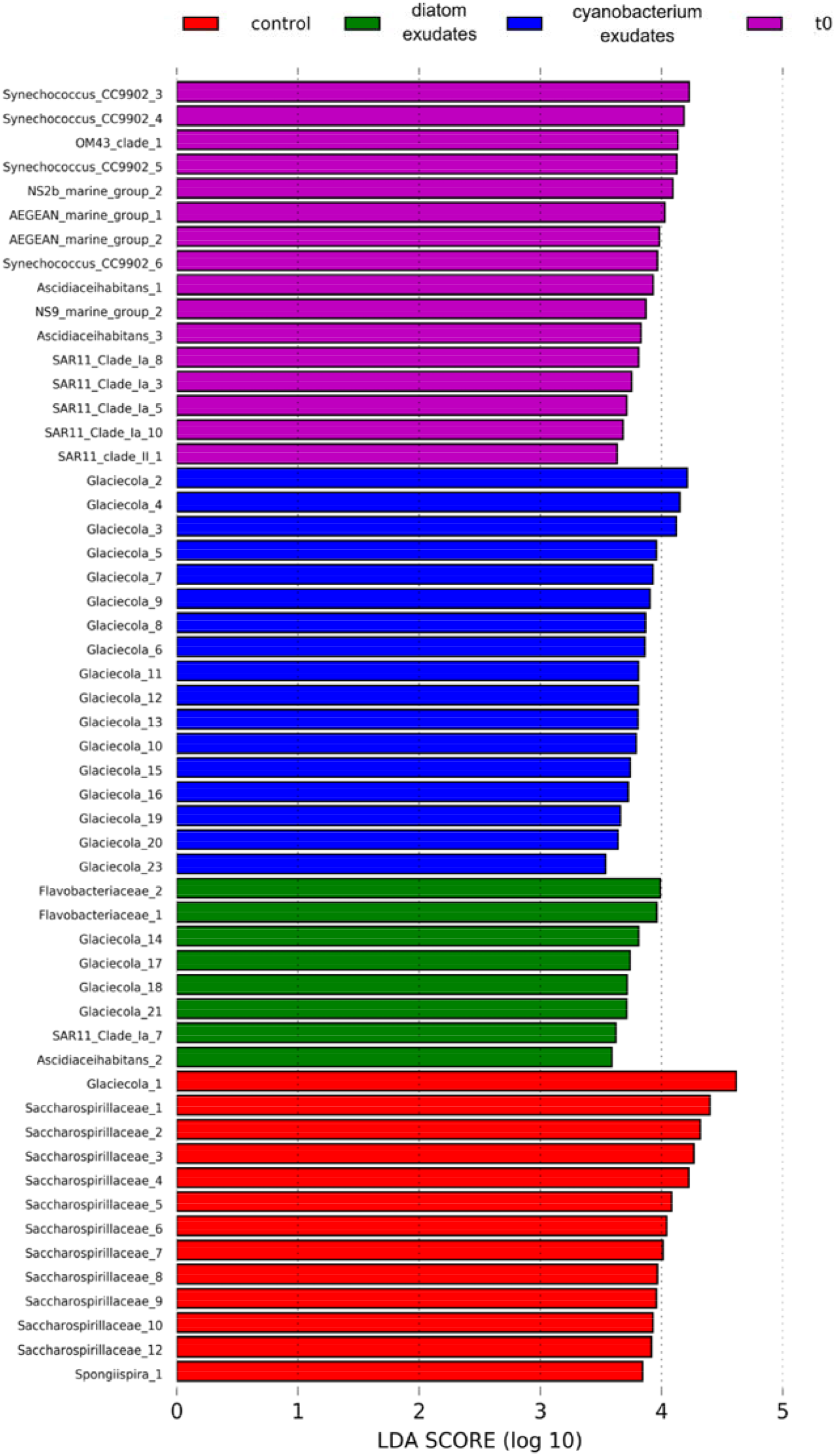
LDA effect size (LEfSe) analyses for definitive ASVs in the treatments t0, control, cyanobacterium exudates and diatom exudates with LDA effect size threshold set to 2.5.

**Suppementary Figure 3B:**
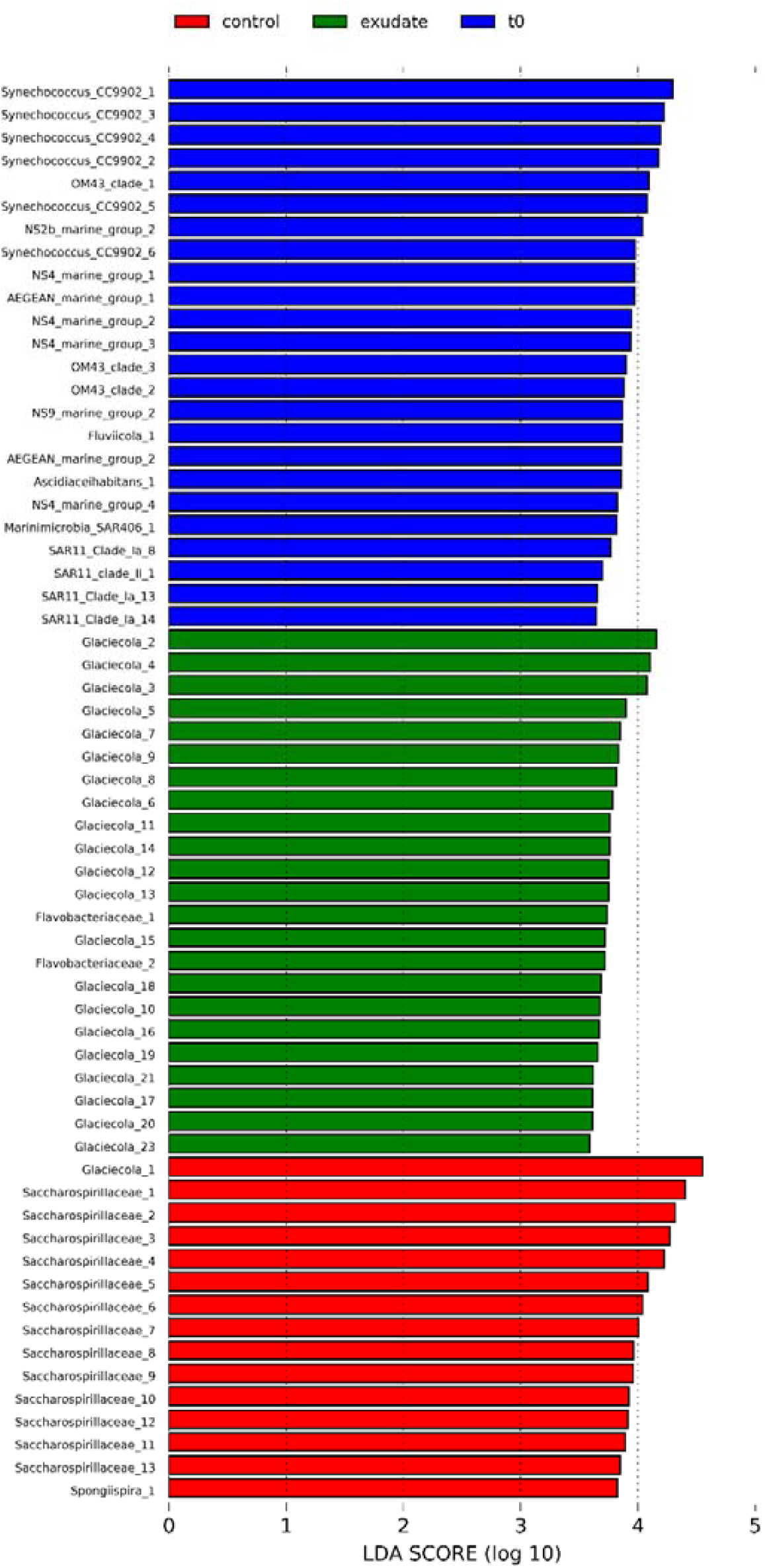
LDA effect size (LEfSe) analyses for definitive ASVs for treatments control, exudates (both phytoplankton donor species), and t0 with LDA effect size threshold set to 2.5.

**Suppementary Figure 3C:**
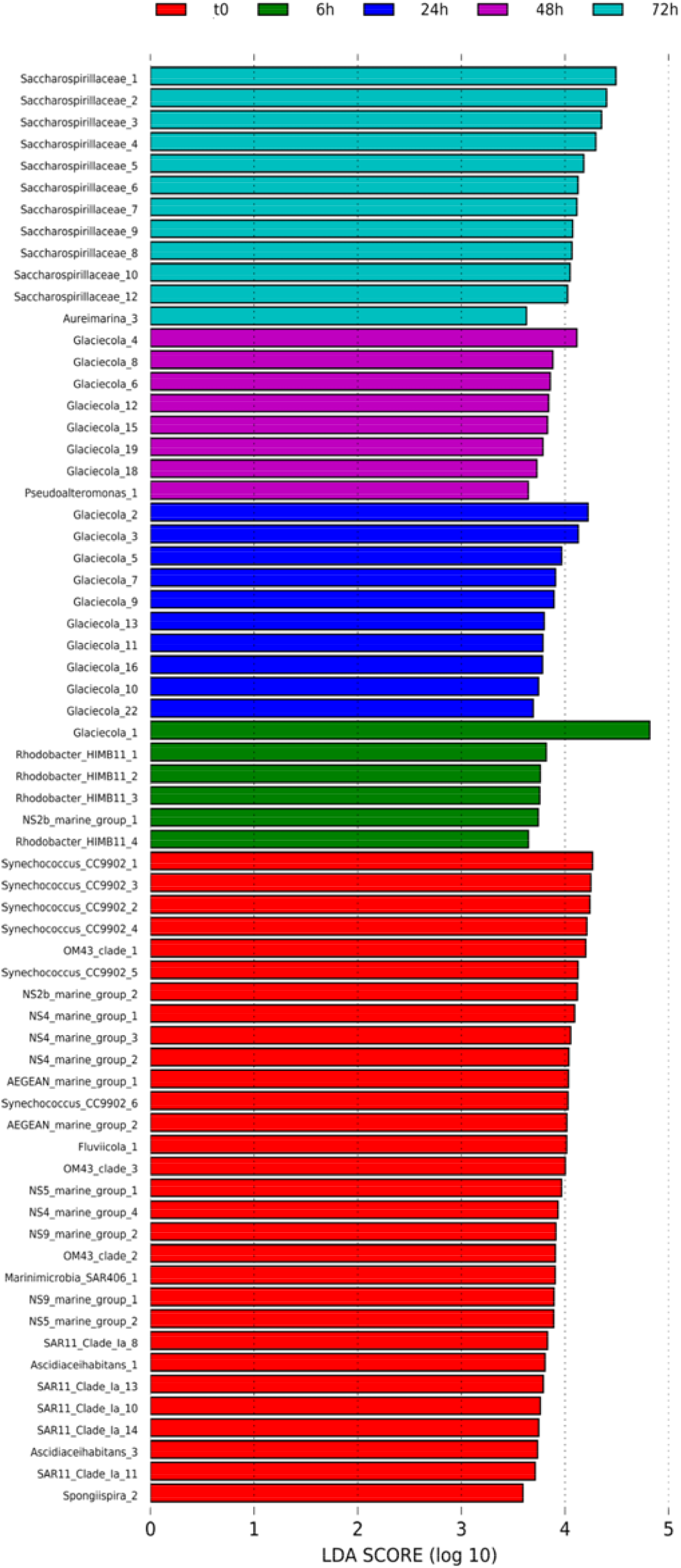
LDA effect size (LEfSe) analyses for definitive ASVs for the different incubation times t0, 6h, 24h, 48h, and 72h with LDA effect size threshold set to 2.5.

**Supplemental Figure 4:**
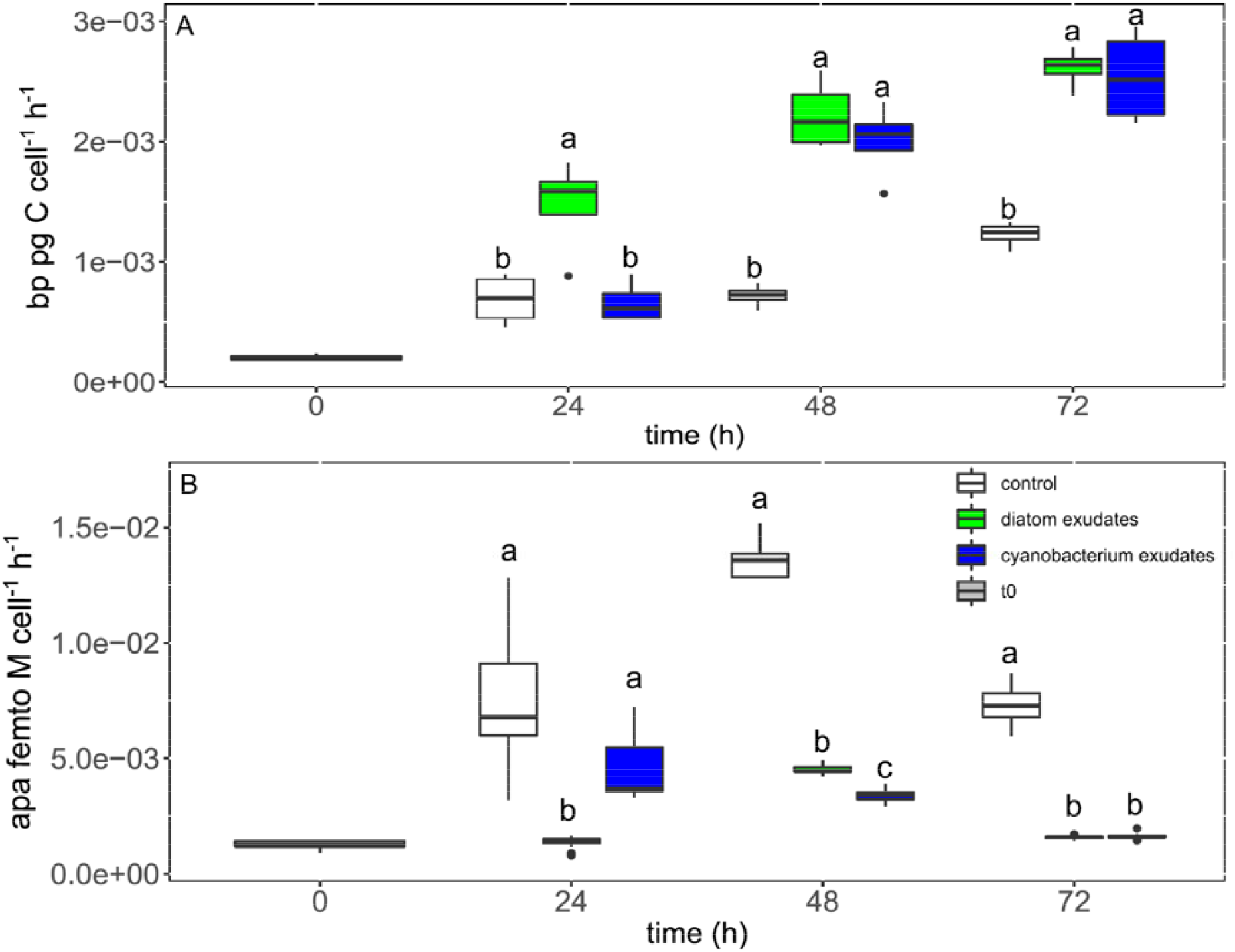
Bacterial production (BP, A) and alkaline phosphatase activity (APA, B)) for the different treatments and time-points calculated on a per cell basis. The letters in the panels represent the outcomes of Tukey post-hoc tests between the treatments for each individual time-point (for further information see Supplemental Table 2).

**Supplemental Table 2:**
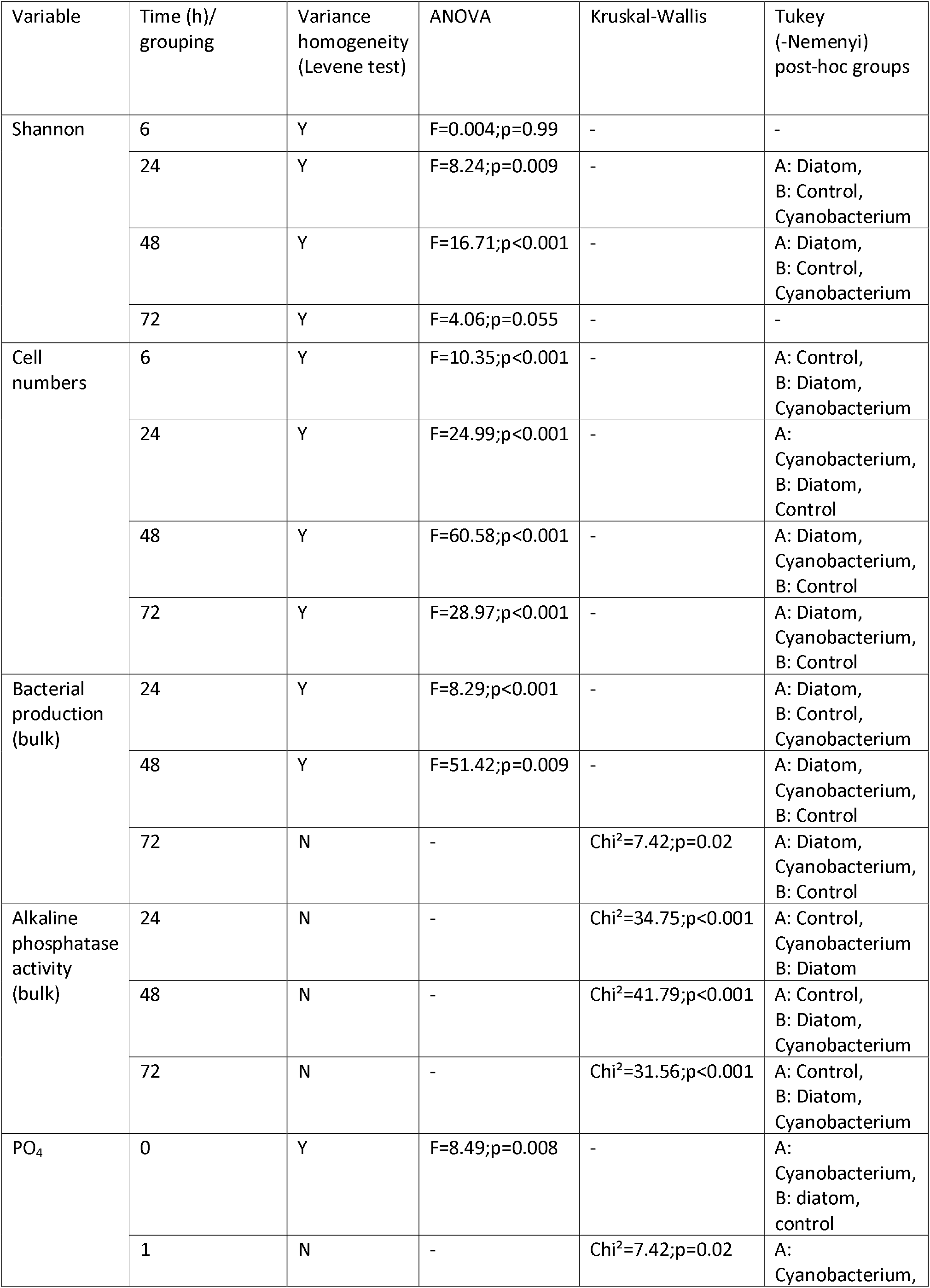
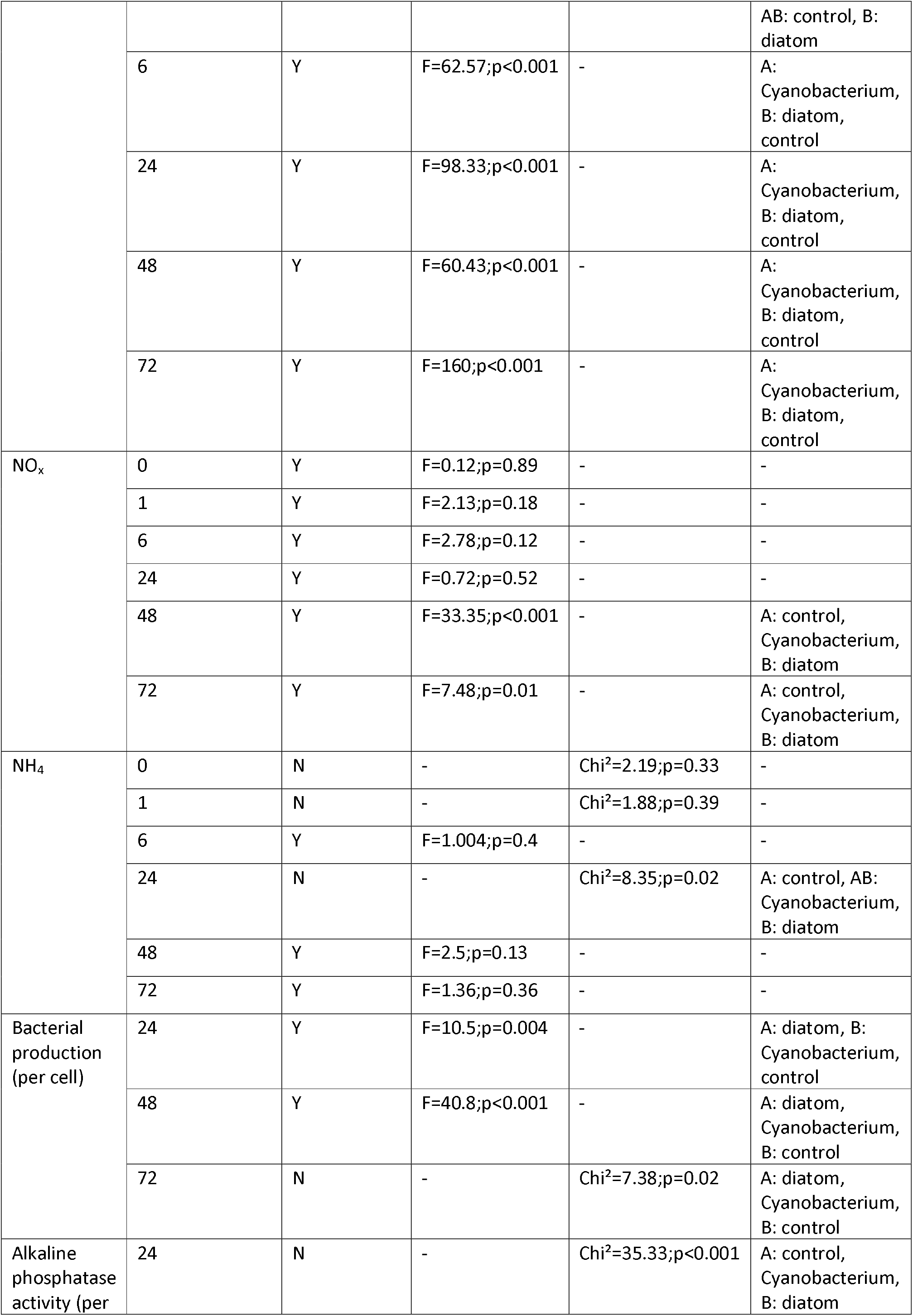
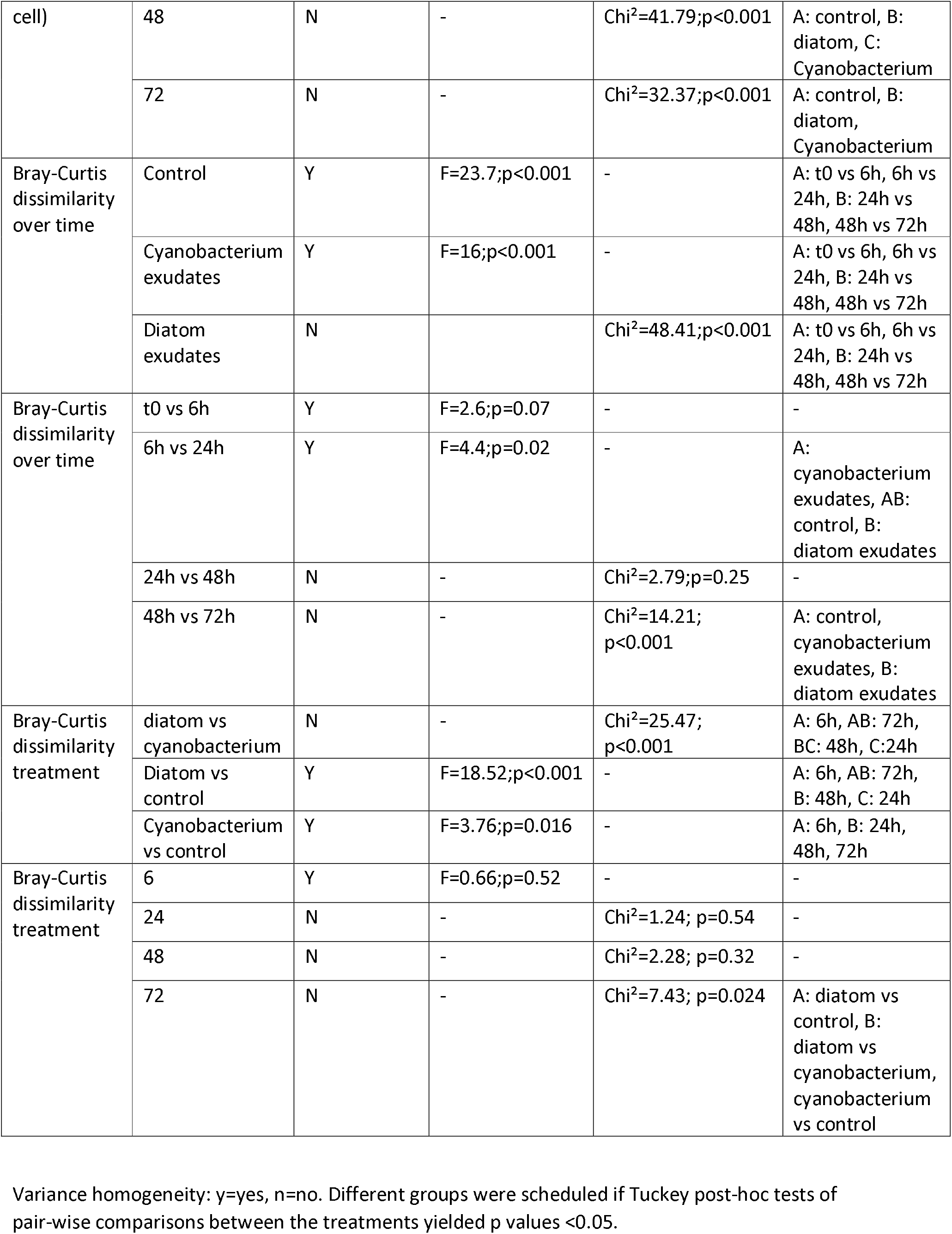
Statistics

